# Delineating bacterial genera based on gene content analysis: a case study of the *Mycoplasmatales*–*Entomoplasmatales* clade within the class *Mollicutes*

**DOI:** 10.1101/2024.08.17.608393

**Authors:** Xiao-Hua Yan, Shen-Chian Pei, Hsi-Ching Yen, Alain Blanchard, Pascal Sirand-Pugnet, Vincent Baby, Gail E. Gasparich, Chih-Horng Kuo

## Abstract

Genome-based analysis allows for large-scale classification of diverse bacteria and has been widely adopted for delineating species. Unfortunately, for higher taxonomic ranks such as genus, establishing a generally accepted approach based on genome analysis is challenging. While core-genome phylogenies depict the evolutionary relationships among species, determining the correspondence between clades and genera may not be straightforward. For genotypic divergence, percentage of conserved proteins (POCP) and genome-wide average amino acid identity (AAI) are commonly used, but often do not provide a clear threshold for classification. In this work, we investigated the utility of global comparisons and data visualization in identifying clusters of species based on their overall gene content, and rationalized that such patterns can be integrated with phylogeny and other information such as phenotypes for improving taxonomy. As a proof of concept, we selected 177 representative genome sequences from the *Mycoplasmatales*–*Entomoplasmatales* clade within the class *Mollicutes* for a case study. We found that the clustering patterns corresponded to the current understanding of these organisms, namely the split into three above-genus groups: Hominis, Pneumoniae, and *Spiroplasma*-*Entomoplasmataceae*-Mycoides (SEM). However, at the genus level, several important issues were found. For example, recent taxonomic revisions that split the Hominis group into three genera and *Entomoplasmataceae* into five genera are problematic, as those newly described or emended genera lack clear differentiations in gene content from one another. Moreover, several cases of mis-classification were identified. These findings demonstrated the utility of this approach and the potential application for other bacteria.

**Impact statement:** Taxonomy provides a foundation for communication that involves biological entities. As such, robust classification and standardized nomenclature are critical. In recent years, genome analysis has been widely adopted for delineating species, but generally accepted approaches for delineating higher taxonomic ranks such as genus are lacking. In this work, we demonstrated that the comparison of overall gene content among species provides an intuitive method for identifying groups of similar organisms that can correspond to genera. Moreover, several critical issues were identified when we applied the method to evaluate recent taxonomic revisions that affected many pathogens with biomedical or economic importance. These findings serve as a cautionary tale against the over-reliance of core-genome centered approaches in taxonomy.

**Data Summary:** The 183 genome assemblies analyzed in this study were obtained from the National Center for Biotechnology Information (NCBI) Genome Database. The accession numbers are provided in Table S1.

## Introduction

Taxonomy involves establishment of hierarchical classification and standardized nomenclature, which provides a foundation for communication about biological entities. For prokaryotic classification at the species level, genome-based approaches are now generally accepted [1–3]. Particularly, methods such as genome-wide average nucleotide identity (ANI) [4, 5] and digital DNA–DNA hybridization (dDDH) [6] are relatively straightforward to implement and interpret, and are therefore widely used. However, for classification at the genus level, establishing a generally accepted approach based on genome analysis is much more challenging [1]. Although similarity indices such as percentage of conserved proteins (POCP) [7] and genome-wide average amino acid identity (AAI) [2, 8] are often used, large variations were observed among different bacteria [9], making the establishment of a universal threshold difficult. As a result, core-genome phylogeny is often used to complement pairwise analysis of genome similarities. However, deciding the correspondence between clades and genera may not be straightforward. As such, even when a reliable phylogeny can be obtained, taxonomic opinions may still vary, particularly between lumpers and splitters. More importantly, core-genome phylogeny reflects only the consensus evolutionary history among a small subset of genes in each individual genome and does not provide any information regarding the gene content diversity, which may have a higher level of relevance to phenotype and ecology. To address this issue, we explore the utility of visualizing global comparisons of gene content divergence among species. If such analysis can provide clear patterns of clustering among species, then the information can be combined with phylogeny, pairwise genome similarities, and other relevant considerations such as phenotypes for more robust classification. As a proof of concept, we conducted a comprehensive analysis for the *Mycoplasmatales*-*Entomoplasmatales* (ME) clade [10] within the class *Mollicutes* [11] that belongs to the phylum *Mycoplasmatota*, formerly named *Tenericutes*.

The class *Mollicutes* includes diverse bacteria that lack a cell wall, have small cell sizes and reduced genomes (usually 0.5-1.5 Mb) with low G+C content (usually 25-30 mol%) [11]. Major groups within this class include ruminant-associated *Anaeroplasma* [12], *Acholeplasma* with diverse habitats [13], plant-pathogenic ‘*Candidatus* Phytoplasma’ that remained uncultivated [14], and the ME clade [10] that contains the majority of described species within *Mollicutes*. Species in the ME clade share an alternative genetic code, in which UGA is assigned to tryptophan instead of being recognized as a stop codon. Notable genera in the ME clade include *Mycoplasma* that contain many important pathogens of human and domestic animals [15], *Spiroplasma* that have helical cell shapes and are mainly arthropod-associated [16], as well as *Entomoplasma* and *Mesoplasma* that evolved from a common ancestor within *Spiroplasma* and lost the helical cell shapes [17]. A major issue in *Mollicutes* taxonomy is that none of these genera (i.e., *Mycoplasma*, *Spiroplasma*, *Entomoplasma*, and *Mesoplasma*) as commonly referred to by relevant communities (e.g., mycoplasmologists, veterinarians, and clinical practitioners) prior to 2018 corresponds to a monophyletic group. For example, the genus *Mycoplasma* is polyphyletic and contains three major groups known as Hominis, Pneumoniae, and Mycoides [15]. Among these, Hominis and Pneumoniae are sister clades and together represent the basal lineage of the ME clade. However, the Mycoides group that includes *Mycoplasma mycoides* subsp. *mycoides*, the type species of the genus, is distantly related to Hominis and Pneumoniae, and evolved from a *Entomoplasma*/*Mesoplasma*-like ancestor after experiencing extensive gene losses and gains [18, 19]. For the other three genera, *Spiroplasma* is a paraphyletic group with multiple clades (i.e., Apis, Citri-Chrysopicola-Mirum, and Ixodetis) [16], while *Entomoplasma* and *Mesoplasma* have Intertwined relationships [17].

Due to this complexity, two proposals for splitting these genera and reclassifying most of the species were made [20, 21]. For the first proposal published in 2018 and focused on *Mycoplasma*, the Hominis group was reclassified into three novel genera (i.e., *Mesomycoplasma*, *Metamycoplasma*, and *Mycoplasmopsis*) and the Pneumoniae group was reclassified into four genera (i.e., *Ureaplasma*, emended *Eperythrozoon*, and newly described *Malacoplasma* and *Mycoplasmoides*) [20]. For the second proposal published in 2019 and focused on *Entomoplasma/Mesoplasma*, both genera were emended and some species were reclassified to newly described *Edwardiiplasma*, *Tullyiplasma*, and *Williamsoniiplasma* [21]. Additionally, these revisions are also associated with changes at higher taxonomic ranks including family and order. With these changes, the aim was to make each of the emended or novel genera correspond to a monophyletic group based on core-genome phylogeny. Additionally, based on the multiple sequence alignments of proteins with diverse functions, a handful of conserved signature indels (CSIs; refers to insertions or deletions in the conserved genes) were identified for each emended or novel genus as the molecular markers for classification. Although these revisions partially solved the issue of non-monophyly, the extensive changes were made without the input of mycoplasmology experts, resulting in debates between the International Committee on Systematics of Prokaryotes (ICSP) Subcommittee on the Taxonomy of Mollicutes and the authors of those works [22–24].

In order to revisit this controversial issue, we analyzed all 177 described species within the ME clade that have genome sequences available. Through improvements in taxon sampling for a global view of genomic divergence within this clade, as well as conducting more detailed characterization within each major group, the aim of this work is to provide additional information related to gene content differentiation to complement approaches such as phylogenetic inference and pairwise comparisons. The findings can provide solid scientific information for further evaluation and discussion regarding the taxonomic revisions by major stakeholders. Moreover, the approach could be adopted for the studies of other prokaryotes.

## Methods

### Data sets

The representative genome assemblies were obtained from the National Center for Biotechnology Information (NCBI) Genome Database. To prioritize those that have passed quality control, the first iteration of our search was conducted based on the NCBI RefSeq data set (ftp://ftp.ncbi.nlm.nih.gov/genomes/ASSEMBLY_REPORTS/assembly_summary_refseq.txt) [25]. Based on prior knowledge regarding *Mollicutes* taxonomy, the ME clade contains three monophyletic groups, namely Hominis, Pneumoniae, and *Spiroplasma*-*Entomoplasmataceae*-Mycoides (SEM) [18, 26]. To include all taxa in these groups, all genus names used before and after recent taxonomic revisions [17, 20, 21] were included to identify the relevant taxa, including *Edwardiiplasma*, *Entomoplasma*, *Eperythrozoon*, *Malacoplasma*, *Mesomycoplasma*, *Mesoplasma*, *Metamycoplasma*, *Mycoplasma*, *Mycoplasmoides*, *Mycoplasmopsis*, *Spiroplasma*, *Tullyiplasma*, *Ureaplasma*, and *Williamsoniiplasma*. Also, relevant taxa with ‘*Candidatus*’ status were included and *Acholeplasma* was selected as the outgroup.

Based on the search conducted on 19 April 2024, 1,651 assemblies belonging to those target genera were identified. Among these, 80 were excluded due to the lack of species-level assignments. From the remaining 1,571 assemblies assigned to 182 species, one assembly was selected as the representative for each species based on the priority rules of: (1) derived from the type strain, (2) identified as the representative of the species by NCBI, and (3) the assembly was complete. To improve taxon sampling, the second iteration of our search was conducted based on the NCBI GenBank data set (ftp://ftp.ncbi.nlm.nih.gov/genomes/ASSEMBLY_REPORTS/assembly_summary_genbank.txt) obtained on the same date to identify species that lack any representative in the RefSeq set. Only one species, *Mycoplasma* (“*Mycoplasmoides*”) *amphoriforme*, was added.

In total, 183 assemblies were included in this study as the “Complete” data set (**Table S1**). Among these, 163 were derived from type strains, including 114 with complete assemblies. For the remaining 20 assemblies that were not derived from type strains, 11 were complete assemblies.

### Comparative and phylogenetic analysis

The procedures of comparative and phylogenetic analysis were largely based on those described in our previous studies [18, 27]. More detailed information is provided in the following sections. Unless stated otherwise, the methods were based on the cited references and the bioinformatics tools were used with the default settings.

For gene content comparison, BLASTP v2.11.0 [28] with e-value cutoff set to 1e^-25^ and OrthoMCL v1.3 [29] were used to infer the homologous gene clusters. For global comparison of gene content divergence, the clustering result was converted into a species-by-gene matrix, with the value in each cell corresponding to the copy number. For visualization, the matrix was plotted as a heatmap using the PHEATMAP package v1.0.12 [30] in R [31]. For principal coordinate analysis (PCoA), the species-by-gene matrix was converted into a Jaccard distance matrix among genomes using the VEGAN package v2.6-4 [32] in R [31], then processed using the “pcoa” function in the APE package v5.8 [33] and visualized using ggplot2 v3.3.2 [34].

For each of the homologous gene clusters that contained single-copy genes conserved in all of the species included in each analysis, multiple sequence alignment was performed using MUSCLE v3.8.31 [35]. The alignment results were manually inspected using Jalview v2.11.3.3 [36]. The concatenated alignments were used to calculate AAI using the PROTDIST function in PHYLIP v3.697 [37] and phylogenetic inference.

For maximum likelihood inference by PhyML v3.3.20180621 [38], the proportion of invariable sites and the gamma distribution parameter were estimated from the data set, the number of substitute rate categories was set to four. The bootstrap support values were estimated based on 1,000 replicates. For validation, Bayesian inference was performed using MrBayes v3.2.7 [39]. The amino acid substitution model was set to mixed with gamma-distributed rate variation across sites and a proportion of invariable sites. The number of rate categories for the gamma distribution was set to four. The Markov chain Monte Carlo analysis was set to run for 1,000,000 generations and sampled every 100 generations, the first 25% of the samples were discarded as the burn-in. To examine the congruence between maximum likelihood and Bayesian inference, the tree topologies were compared using the TREEDIST function in PHYLIP v3.697 [37] with the “symmetric difference” option. For visualization of the phylogenetic trees, the Interactive Tree of Life (iTOL) v6 [40] was used.

## Results

### Phylogeny

Among the 183 genomes analyzed, 30 single-copy genes were conserved. Phylogenetic inference based on the concatenated alignment using the maximum likelihood and Bayesian methods produced highly congruent results with strong support (**Figure S1**). Importantly, the placements of those 177 ingroup species into the three groups defined previously (i.e., SEM, Hominis, and Pneumoniae) were entirely consistent between the two methods. The relationships among these three groups were also consistent with our current understanding of the ME clade [17–19]. For additional validation, a more conservative approach for homolog identification and multiple sequence alignment was used prior to maximum likelihood inference. Based on a set of 23 single-copy genes and trimmed alignment of these genes, a phylogeny that produced the same placement of those 177 ingroup species into the three groups was obtained (**Figure S2**).

To increase the number of conserved genes for more robust inference and to improve visualization, we conducted the further phylogenetic inference for a “Selected” data set with 47 representatives (**Figure 1**) and each of the three groups separately (**Figure S3-5**). For the SEM group (**Figures 1 and S3**), the phylogeny was consistent with previous works by us [17, 18] and Gupta et al. [21], although the genus assignments were different due to our differences in taxonomic opinions.

**Figure 1.**
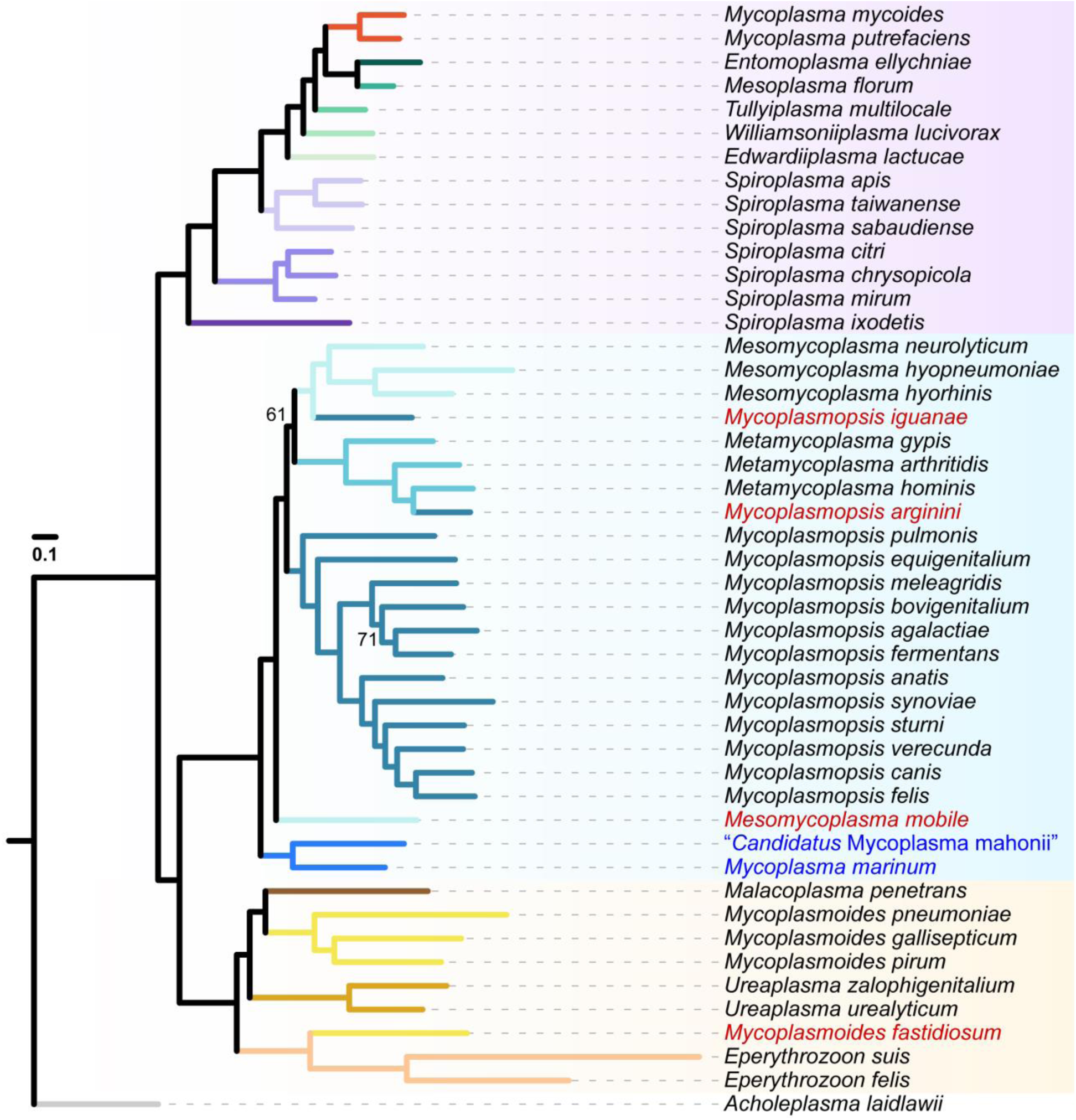
Molecular phylogeny of the *Mycoplasmatales*-*Entomoplasmatales* clade. This maximum likelihood phylogeny included 47 selected species and was based on a concatenated alignment of 75 conserved single-copy genes with 37,172 aligned amino acid sites. Branches were color-coded according to the taxonomic assignments. *Acholeplasma laidlawii* was included as the outgroup. The four species with names highlighted in red had phylogenetic placements inconsistent with expectation based on the recent taxonomic revisions. The two species highlighted in blue represent a novel branch that was not included in the recent taxonomic revisions. The bootstrap support was estimated based on 1,000 resampling. Only two nodes had a bootstrap value < 80% and the exact values were labeled. A Bayesian inference based on the same alignment produced an identical topology and all nodes have > 90% posterior probability support.

For the Hominis group (**Figures 1, S1, and S4**), several major issues were found. First, three newly described species isolated from marine invertebrates (i.e., *Mycoplasma marinum*, *Mycoplasma totarodis*, and “*Candidatus* Mycoplasma mahonia”) [41, 42] formed a novel sub-clade distinct from any of the three newly described genera (i.e., *Mesomycoplasma*, *Metamycoplasma*, and *Mycoplasmopsis*). Second, 14 other species, including 12 described after 2018 (i.e., *Mycoplasma anserisalpingitidis*, *Mycoplasma enhydrae*, *Mycoplasma miroungigenitalium*, *Mycoplasma miroungirhinis*, *Mycoplasma nasistruthionis*, *Mycoplasma phocimorsus*, *Mycoplasma phocoenae*, *Mycoplasma phocoeninasale*, *Mycoplasma procyoni*, *Mycoplasma seminis*, *Mycoplasma struthionis*, and *Mycoplasma tauri*) and two that had genome sequences available in 2021 (i.e., *Mycoplasma zalophi* and *Mycoplasma zalophidermidis*), did not adopt the revised taxonomy and are still classified as members of *Mycoplasma*, such that all three newly described genera for this clade became non-monophyletic. Third, three species were found to have phylogenetic placements that conflict with the revised taxonomy [20]: (1) *Mycoplasmopsis iguanae* was classified based on only the 16S rRNA gene [20], but the core-genome phylogenies indicated that it is more closely related to *Mesomycoplasma*; (2) *Mycoplasmopsis arginini* was classified based on a problematic assembly that has been removed from NCBI RefSeq (GCF_000428625.1), and our updated analysis based on another assembly (GCF_900660725.1) indicated this species should be a member of *Metamycoplasma*; and (3) *Mesomycoplasma mobile* had unstable phylogenetic placements in the previous work [20], and our updated analysis indicated that it does not belong to any of the three newly described genera based on the criterion of monophyly.

Due to these issues, we examined the 12 CSIs used to define those three novel genera within the Hominis clade. We found that in addition to the seven exceptions reported previously [20], 45 additional inconsistencies were found, including nine cases that involve species with genome sequences available before 2018 and 36 cases that involve species with genome sequences available after 2018 (**Table S2**). In total, 52 cases of exceptions or inconsistencies were found and affect all of the 12 CSIs. These findings raised concerns regarding the reliability of using these CSIs for classification.

For the Pneumoniae group, the relationships among the four genera differed between the complete data set (**Figures S1 and S5**) and the “Selected” data set (**Figure 1**). The incongruence may be caused by the long branches within this group, such that more comprehensive taxon sampling is important. The results of the complete data set, either when analyzed together with other groups (**Figure S1**) or restricted to the Pneumoniae group (**Figure S5**), may be more reliable as this tree topology is also consistent with the previous work [20]. However, one species that was classified based on only the 16S rRNA gene (i.e., *Mycoplasmoides fastidiosum*) [20] would need to be re-classified if the monophyly of this genus is to be maintained. Additionally, three other species are still classified under the genus *Mycoplasma* (i.e., *Mycoplasma bradburyae*, *Mycoplasma tullyi*, and “*Candidatus* Mycoplasma haemohominis”) (**Figure S5**).

### Genome similarity indices

To investigate if genome similarity indices such as POCP [7] or AAI [2, 8] can be used for delineating genus within the ME clade, we performed all pairwise comparisons among the genomes analyzed. A high level of variation in the distribution of these values was observed among the three major groups (**Figure S6**). Data visualization based on the “Selected” data set also indicated that no universal threshold can be established for differentiating different genera among these groups (**Figure 2**).

**Figure 2.**
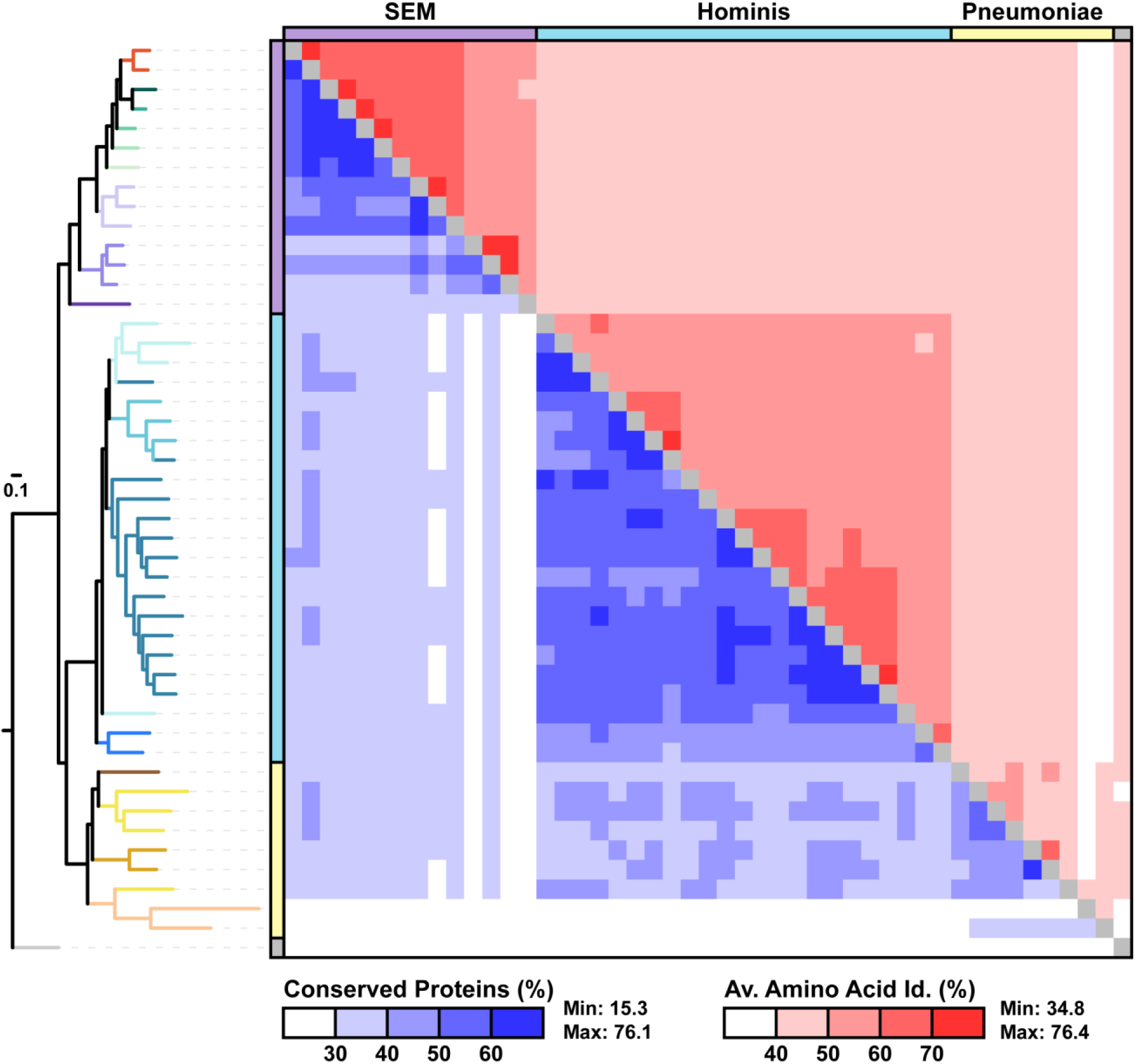
Genome similarities among representative *Mollicutes* species. The tree was based on the maximum likelihood phylogeny shown in Figure 1. Two measurements of pairwise genome similarities were used, including the percentage of conserved proteins (POCP) (below the diagonal) and the average amino acid identity (AAI) of those conserved proteins (above the diagonal).

### Gene content

To explore the utility of gene content comparisons in classification, we inferred the pan-genome for visualization. In total, 32,496 homologous gene clusters were identified. Visualization of the 3,637 homologous gene clusters shared by at least five species illustrated that species that are more closely related based on core-genome phylogeny also tend to be more similar in gene content (**Figure 3**). However, the similarity pattern is quite noisy, which is also expected based on the extensive gene gains and losses in these bacteria [18, 19, 43–45]. As such, identification of genus-specific genes as molecular markers for classification is difficult and may not be a robust approach as more genome sequences become available.

**Figure 3.**
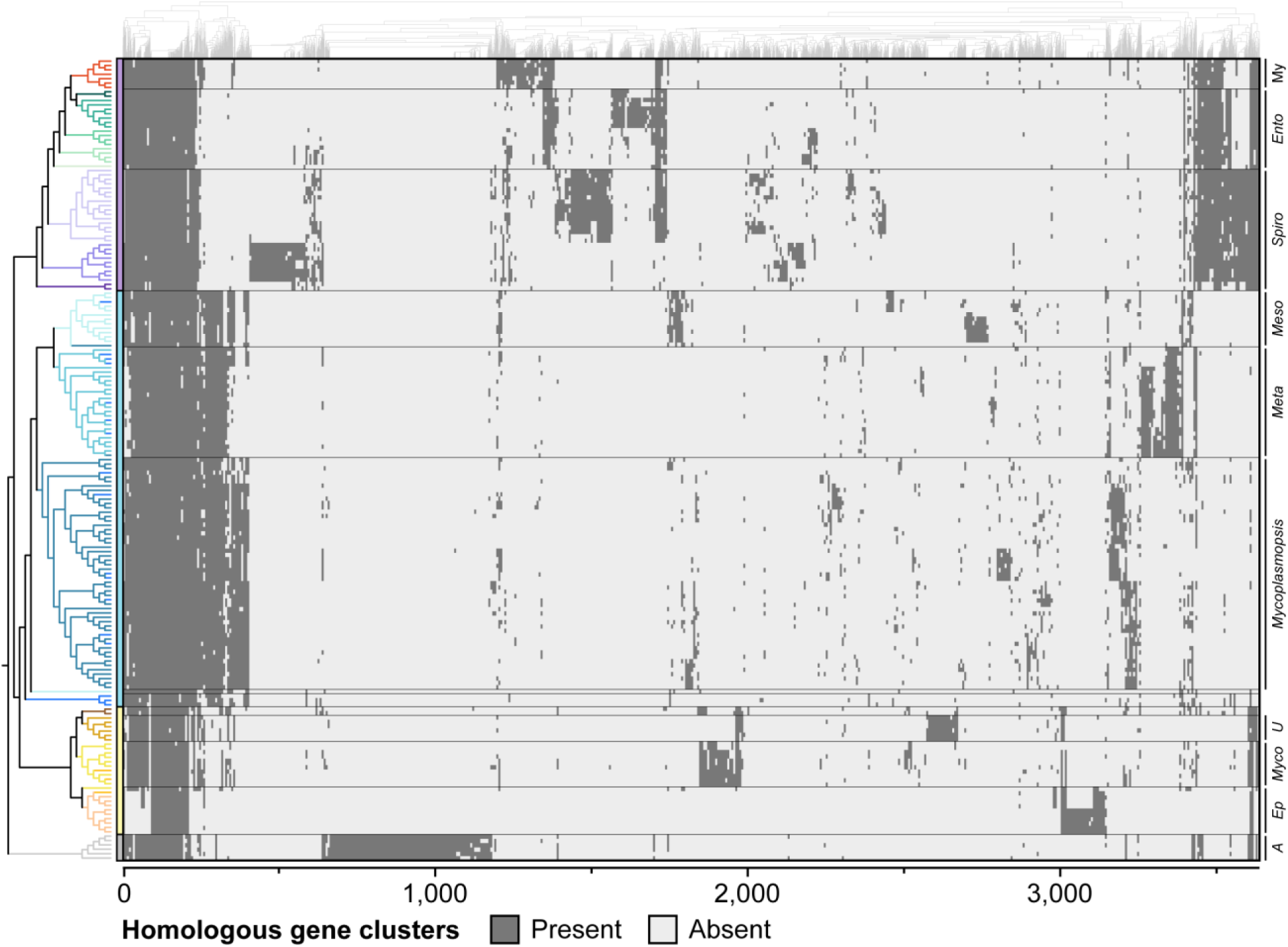
Gene content heatmap. The rows correspond to species and the columns correspond to homologous gene clusters. The cladogram containing all 183 species in the complete data set was based on the maximum likelihood phylogeny shown in Figure S1. The columns were arranged based on hierarchical clustering of distribution patterns. Among of the 32,496 homologous gene clusters identified in the pan-genome of these species, only 3,637 clusters that contain homologs found in at least five species were included in the visualization.

For a simplified and more intuitive visualization of overall gene content divergence, we conducted PCoA and plotted the results based on the first two coordinates (**Figure 4**). In these plots, each dot represents one species and the distance between dots corresponds to their divergence. As such, if clear clustering could be observed among the species analyzed, each cluster could correspond to a taxonomic unit defined by similarity in overall gene content, and the area encompassed by a cluster of species would correspond to their gene content diversity. When the 177 ingroup species were analyzed together (**Figure 4a**), a clustering pattern corresponding to the three major groups (i.e., SEM, Hominis, and Pneumoniae) could be observed. However, correspondence between the clustering pattern within each of the three major groups and the current taxonomy was more complicated. For the SEM group, the three sub-groups of *Spiroplasma* (i.e., Apis, Citri-Chrysopicola-Mirum, and Ixodetis) and the Mycoides *Mycoplasma* all formed well-separated clusters, with species within the same cluster being more similar to each other while divergent from those in other clusters. However, the 18 species belonging to *Entomoplasma*, *Mesoplasma*, *Edwardiiplasma*, *Tullyiplasma*, and *Williamsoniiplasma* did not form five distinct clusters. Rather, those 18 species formed one single cluster distinct from all *Spiroplasma* clusters or the Mycoides *Mycoplasma* cluster. The same pattern could be observed when the analysis included only the 53 SEM group species (**Figure 4b**).

**Figure 4.**
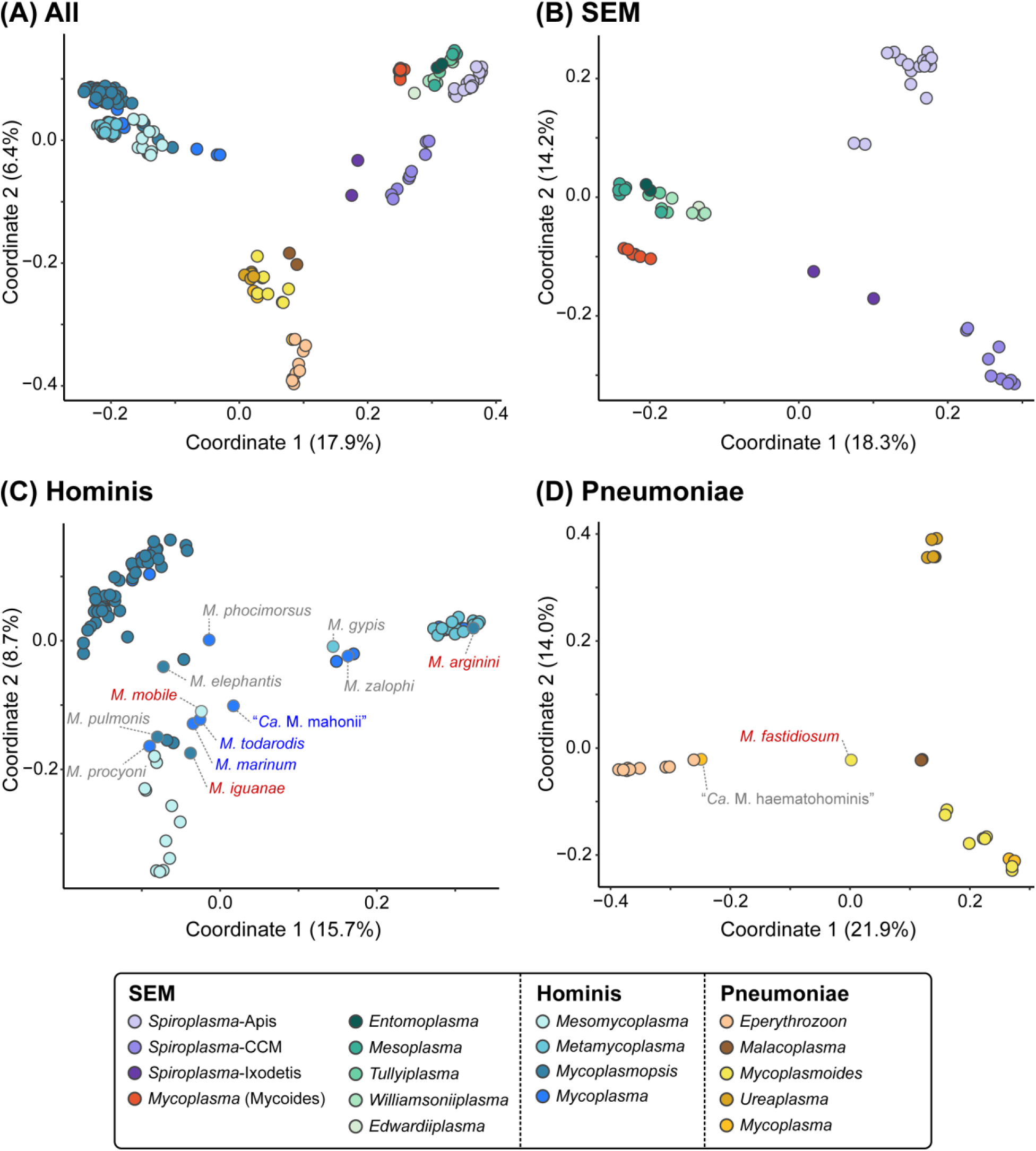
Principle coordinate analysis of gene content differentiation. Dots representing individual species are color-coded according to the genus/clade assignment. Distance between dots indicates the level of dissimilarity. The percentage of variance explained by each axis is provided in parentheses. (A) All three major groups within the *Mycoplasmatales*-*Entomoplasmatales* clade. (B) The 53 species within the *Spiroplasma*-*Entomoplasmataceae*-Mycoides (SEM) group. (C) The 95 species within the Hominis group. (D) The 29 species within the Pneumoniae group.

For the Hominis group, while clustering could be observed for *Mesomycoplasma* and *Metamycoplasma*, both of these genera were encompassed within the diversity of *Mycoplasmopsis* (**Figure 4a**). This finding indicated that when the gene content diversity of the entire ME clade was considered as a whole, species belonging to the Hominis group may be considered as a relatively homogeneous group without robust boundaries for delineation into different genera. When the analysis included only those 95 Hominis group species, a higher resolution for distinguishing the gene content divergence among the newly described genera may be obtained (**Figure 4c**). Species in the genus *Metamycoplasma* species formed two clusters, which is congruent with the two sub-clades in the core-genome phylogeny (**Figure S4**). However, the separation of *Mesomycoplasma* and *Mycoplasmopsis* species was not so clear cut, and those basal lineages (i.e., *M. mobile*, *M. marinum*, *M. todarodis*, and “*Ca*. M. mahonii”) appeared to be more similar to *Mycoplasmopsis* species in their gene content.

For the Pneumoniae group, an overlap between *Mycoplasmoides* and *Ureaplasma* was observed when all ME clade species were analyzed jointly (**Figure 4a**). When the analysis included only the 29 Pneumoniae group species, species belonging to different genera became better separated (**Figure 4d**) and the pattern was congruent with expectations based on phylogeny (**Figure S5**). Notably, *Mycoplasmoides fastidiosum* appeared to be divergent from any of the four genera described within this clade (i.e., *Eperythrozoon*, *Malacoplasma*, *Mycoplasmoides*, and *Ureaplasma*). This finding, together with its phylogenetic placement (**Figures 1, S1, and S5**), suggested that this species represents a distinct lineage within the Pneumoniae clade both in terms of gene content and core-genome sequence divergence, and the previous assignment of this species to *Mycoplasmoides* should be emended.

## Discussion

In taxonomy, robust classification of organisms into well-defined groups requires polyphasic approaches. Unfortunately, selecting the phenotypic traits for consideration often requires domain knowledge from experts of specific taxonomic groups, and it can be difficult to weigh different traits objectively. Consequently, phenotypic approaches often play a minor role or are even ignored in large-scale studies. In contrast, with the improved availability of genome sequences and computation power over the past decade, core-genome phylogeny has started to gain popularity in taxonomic studies, particularly for large-scale analysis [46, 47]. However, overemphasizing the core-genome phylogeny can result in several problems. First, when only a small number of core genes are included, which is often the case when the analysis involves highly divergent and/or genome-reduced organisms such as the class *Mollicutes* in this study, the phylogeny may not be reliable. Second, even when a robust phylogeny can be inferred, the resulting tree reflects only the combined evolutionary history of those core genes, which account for only a small proportion of all genes in any of the organisms studied. In other words, core-genome phylogeny does not necessarily reflect the evolutionary histories of other genes in the genome. Third, due to rate heterogeneity among different organisms, objectively determining a branch length to define a given taxonomic rank (e.g., genus) is difficult, and often leads to arguments between lumping and splitting with no resolution.

In this work and our previous studies on different bacteria [18, 27, 48, 49], we demonstrated that the comparisons of overall gene content could be a useful approach for evaluating genotypic divergence at different taxonomic ranks, which supplements results from core-genome phylogeny. Particularly, PCoA provides a simple and intuitive method for data visualization. If clear patterns of clustering could be observed, organisms belonging to the same cluster would be similar to each other in their gene content, while different from those in other clusters. Given the plausible link between overall gene content and other aspects of organismal biology, the clustering patterns observed in PCoA plots could facilitate robust classification. While this approach cannot completely resolve possible debates between lumping and splitting, as the breadth of taxon sampling would affect the clustering patterns, comparing the results obtained from different sampling could still provide useful information.

For example, while whether *Metamycoplasma* species could be clearly distinguished from other genera depends on if the analysis included the entire ME clade (**Figure 4a**) or only the Hominis clade (**Figure 4c**), in both analyses the boundaries between *Mesomycoplasma* and *Mycoplasmopsis* remained unclear. This observation, together with the short internal branches that separate those newly described genera (**Figures 1 and S1**), raised the concern regarding the robustness of splitting the Hominis clade species into those three genera [20]. Furthermore, with the discovery of the novel clade represented by *M. marinum* and the classification of all these species to the genus *Mycoplasma* [41, 42], the issue of *Mycoplasma* being a polyphyletic group remains unresolved.

In another example concerning the SEM clade, the gene content comparison approach provided more robust and clear patterns. For those 18 *Entomoplasmataceae* species, two proposals for taxonomic revisions were published in 2019. While Gupta et al. argued for splitting these species into five genera (i.e., emended *Entomoplasma* and *Mesoplasma*, together with the newly described *Edwardiiplasma*, *Tullyiplasma*, and *Williamsoniiplasma*) [21], we suggested lumping all 18 species into one single genus (i.e., emended *Entomoplasma*) [17]. Considering the shared characteristics among these 18 species in comparison with other SEM species such as cell morphology, metabolism, and ecology [17, 18], the robustness of gene content comparisons results across different sampling breadth (**Figure 4 panels A and B**) as presented in this work provided further support for lumping.

Regarding the possible pitfalls of this gene content comparison approach, data visualization based on only the first two coordinates with the highest percentages of variance explained could result in considerable signal loss. In the four PCoA plots shown in this work (**Figure 4**), the first two coordinates explained 24.3% to 35.9% of the variance when combined. While this is non-ideal, such signal loss is inevitable for statistical methods that involve dimensionality reduction. With pan-genome size reaching tens of thousands of homologous genes and each homologous gene represents one dimension in quantifying the genotypic divergence among genomes, such signal loss should be a realistic expectation and acceptable. In comparison, simple pairwise comparisons such as PCOP or AAI that compress the overall divergence between genomes into a single number likely resulted in even higher levels of signal loss, and often produce continuous distributions of similarity values that do not have a clear cutoff for defining taxonomic groups (**Figures 2 and S6**). Of course, including genome assemblies that are incomplete or have low quality annotation in the analysis could bias the result due to false gene absence, and such cases should be treated carefully. In this work, the emphasis of using NCBI RefSeq data sets and complete assemblies were both considerations against such potential pitfalls.

Regarding the specific organisms analyzed in this study, multiple critical issues were identified for the recent taxonomic revisions [20, 21]. First, several mistakes were found, including problematic classification of those four species as described in the Results section (**Figures 1 and S1**). The mis-classifications based on only the 16S rRNA gene sequences were unsurprising but still problematic. There are 20 other species that were re-classified based on only the 16S rRNA gene sequences, but still do not have any genome sequence available. Future genomic characterization of these species may uncover more cases of mis-classifications. Second, the reliance of CSIs as molecular markers for classification is highly questionable. There is no guarantee that such sequence polymorphisms could be reliable synapomorphies, as horizontal gene transfer or convergent evolution could disrupt the pattern. In practice, as shown in our re-examination of the Hominis clade, improved availability of genome sequences from different lineages greatly increased the number of exceptions to the established rules for classification (**Table S2**). As more novel species or strains are discovered in the future, the system may eventually become impractical.

## Conclusion

Regarding the question of “How to define a genus in bacteria?”, there may not be a universal answer applicable across different groups. Nevertheless, the gene content comparison approach as presented in this work, together with core-genome phylogeny, provide useful information for evaluating and improving taxonomy. The approach of lumping versus splitting each has its pros and cons. For example, prior to the 2018 revision [20], the genus *Mycoplasma* included species belonging to different groups (i.e., Hominis, Pneumoniae, Mycoides). While this taxonomic treatment has its historical reasons and has been widely adopted, the extensive diversity and the polyphyly of this group were considered as issues that justified the taxonomic revision [23]. On the other hand, the 2018 revision [20] kept only seven species in the emended *Mycoplasma* and reclassified > 100 species to five novel genera. While the splitting made each of the emended and newly described genera corresponds to a monophyletic group based on the core-genome phylogeny, which was rationalized as improvements that can facilitate communication [23], the sweeping changes and the creation of many new names can cause confusion that affects research, diagnosis, therapy, and legislation [22]. Worse, as demonstrated in this work, the revision contained multiple errors that made those newly described genera non-monophyletic (**Figure 1**), the proposed molecular markers for classification were unreliable (**Table S2**), and some of the splitting appeared to be questionable (**Figure 4**). Considering that the new nomenclature has been adopted by major databases, including the NCBI Taxonomy [25] and the List of Prokaryotic names with Standing in Nomenclature (LPSN) [50], as well as some scientific publications, these errors and issues are detrimental to communication. A lesson learned from this case is that for extensive taxonomic revisions, particularly those affecting organisms with biomedical or economic importance, the proposals should be examined carefully by experts with appropriate domain knowledge, and potentially premature revisions should be adopted with extreme caution. Importantly, for proper evaluation of taxonomic revisions, investigations relying on only core-genome phylogeny or simple pairwise comparisons (e.g., PCOP and AAI) do not provide sufficient information. The gene content comparison approach described in this study can provide useful information on genotypic divergence to facilitate the evaluation of taxonomic revisions for improvements.

## Author Contributions

Conceptualization: AB, CHK

Funding acquisition: CHK

Investigation: XHY, SCP, HCY, AB, VB, CHK

Methodology: PSP, VB, CHK

Project administration: CHK

Resources: AB, PSP, VB, CHK

Supervision: AB, CHK

Validation: XHY, SCP, HCY, AB

Visualization: XHY, SCP, HCY

Writing – original draft: CHK

Writing – review & editing: AB, GEG, CHK

## Acknowledgements

We thank members of the International Organization for Mycoplasmology (IOM) and the Subcommittee on the Taxonomy of Mollicutes under the International Committee on Systematics of Prokaryotes (ICSP) for scientific discussions that inspired this work. This study benefited from Calcul Québec (https://www.calculquebec.ca/) and l’Alliance de Recherche Numérique du Canada (https://www.alliancecan.ca/) for the access to the computer expertise and time required for some of the phylogenetic analyses.

## Supplementary Materials

**Table S1. List of the genome assemblies used in this study.**

**Table S2. List of inconsistencies found for the conserved signature indels (CSIs).**

**Figure S1. Molecular phylogeny of the *Mycoplasmatales*-*Entomoplasmatales* clade.**

**Figure S2. Comparison of maximum likelihood trees produced using different approaches.**

**Figure S3. Molecular phylogeny of the *Spiroplasma*-*Entomoplasmataceae*-Mycoides (SEM) group.**

**Figure S4. Molecular phylogeny of the Hominis group.**

**Figure S5. Molecular phylogeny of the Pneumoniae group.**

**Figure S6. Frequency distribution of pairwise genome similarities.**

