## Supplementary material for "Delineating bacterial genera based on gene content analysis: a case study of the *Mycoplasmatales*–*Entomoplasmatales* clade within the class *Mollicutes*": Figure S1

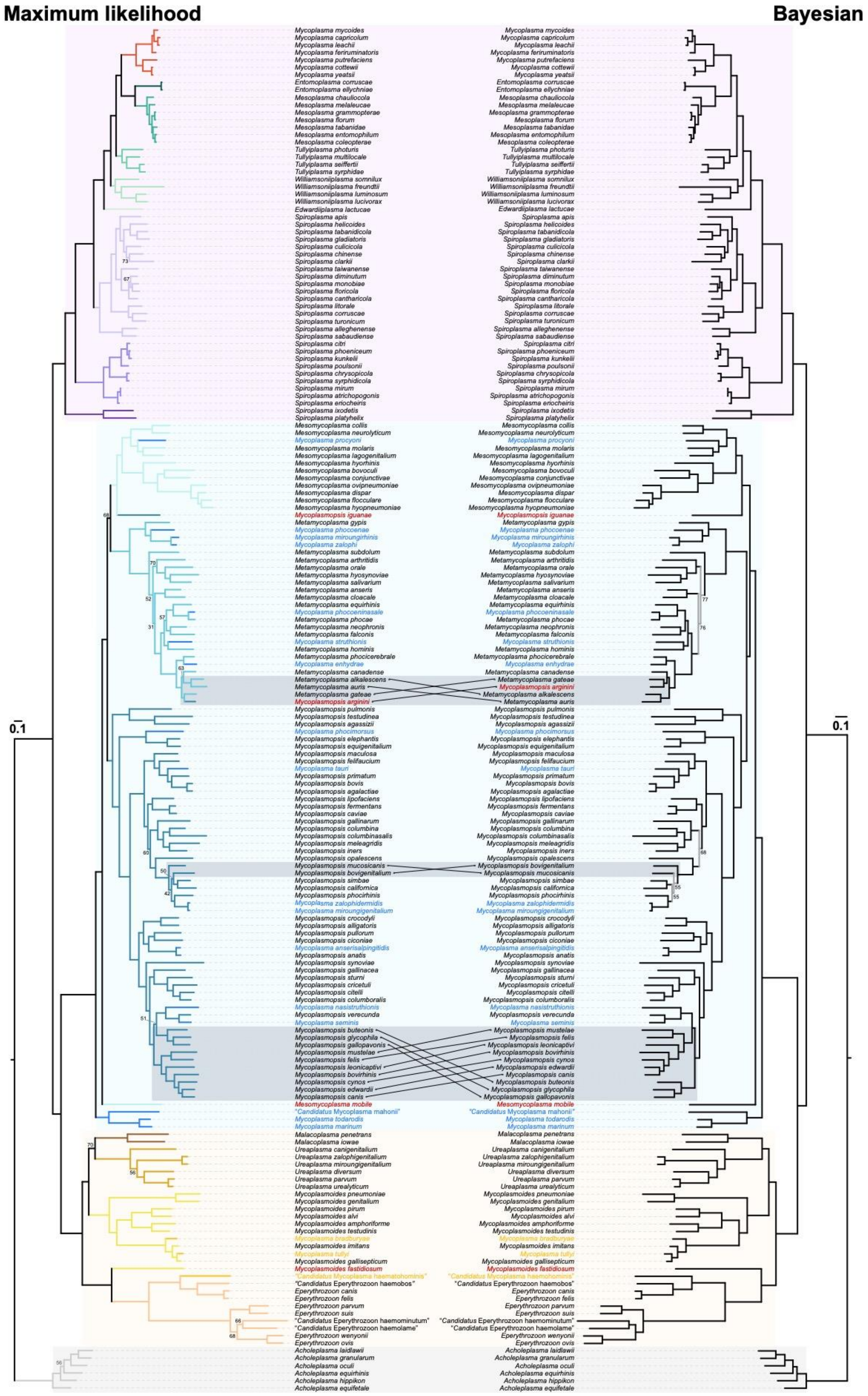
Maximum likelihood
Bayesian
0.1
0.1
Mycoplasma mycoides
Mycoplasma capricolum
Mycoplasma leachii
Mycoplasma feriruminatoris
Mycoplasma putrefaciens
Mycoplasma cottewii
Mycoplasma yeatsii
Entomoplasma corruscae
Entomoplasma ellychniae
Mesoplasma chauliocola
Mesoplasma melaleucae
Mesoplasma grammopterae
Mesoplasma florum
Mesoplasma tabanidae
Mesoplasma entomophilum
Mesoplasma coleopterae
Tullyiplasma photuris
Tullyiplasma multilocale
Tullyiplasma seiffertii
Tullyiplasma syrphidae
Williamsoniiplasma somnilux
Williamsoniiplasma freundtii
Williamsoniiplasma luminosum
Williamsoniiplasma lucivorax
Edwardiiplasma lactucae
Spiroplasma apis
Spiroplasma helicoides
Spiroplasma tabanidicola
Spiroplasma gladiatoris
Spiroplasma culicicola
Spiroplasma chinense
Spiroplasma clarkii
Spiroplasma taiwanense
Spiroplasma diminutum
Spiroplasma monobiae
Spiroplasma floricola
Spiroplasma cantharicola
Spiroplasma litorale
Spiroplasma corruscae
Spiroplasma turonicum
Spiroplasma alleghenense
Spiroplasma sabaudiense
Spiroplasma citri
Spiroplasma phoeniceum
Spiroplasma kunkelii
Spiroplasma poulsonii
Spiroplasma chrysopicola
Spiroplasma syrphidicola
Spiroplasma mirum
Spiroplasma atrichopogonis
Spiroplasma eriocheiris
Spiroplasma ixodetis
Spiroplasma platyhelix
Mesomycoplasma collis
Mesomycoplasma neurolyticum
Mycoplasma procyoni
Mesomycoplasma molaris
Mesomycoplasma lagogenitalium
Mesomycoplasma hyorhinis
Mesomycoplasma bovoculi
Mesomycoplasma conjunctivae
Mesomycoplasma ovipneumoniae
Mesomycoplasma dispar
Mesomycoplasma flocculare
Mesomycoplasma hyopneumoniae
Mycoplasmopsis iguanae
Metamycoplasma gypis
Mycoplasma phocoenae
Mycoplasma miroungirhinis
Mycoplasma zalophi
Metamycoplasma subdolum
Metamycoplasma arthritidis
Metamycoplasma orale
Metamycoplasma hyosynoviae
Metamycoplasma salivarium
Metamycoplasma anseris
Metamycoplasma cloacale
Metamycoplasma equirhinis
Mycoplasma phocoeninasale
Metamycoplasma phocae
Metamycoplasma neophronis
Metamycoplasma falconis
Mycoplasma struthionis
Metamycoplasma hominis
Metamycoplasma phocicerebrale
Mycoplasma enhydrae
Metamycoplasma canadense
Metamycoplasma alkalescens
Metamycoplasma auris
Metamycoplasma gateae
Mycoplasmopsis arginini
Mycoplasmopsis pulmonis
Mycoplasmopsis testudinea
Mycoplasmopsis agassizii
Mycoplasma phocimorsus
Mycoplasmopsis elephantis
Mycoplasmopsis equigenitalium
Mycoplasmopsis maculosa
Mycoplasmopsis felifaucium
Mycoplasma tauri
Mycoplasmopsis primatum
Mycoplasmopsis bovis
Mycoplasmopsis agalactiae
Mycoplasmopsis lipofaciens
Mycoplasmopsis fermentans
Mycoplasmopsis caviae
Mycoplasmopsis gallinarum
Mycoplasmopsis columbina
Mycoplasmopsis columbinasalis
Mycoplasmopsis meleagridis
Mycoplasmopsis iners
Mycoplasmopsis opalescens
Mycoplasmopsis mucosicanis
Mycoplasmopsis bovigenitalium
Mycoplasmopsis simbae
Mycoplasmopsis californica
Mycoplasmopsis phocirhinis
Mycoplasma zalophidermidis
Mycoplasma miroungigenitalium
Mycoplasmopsis crocodyli
Mycoplasmopsis alligatoris
Mycoplasmopsis pullorum
Mycoplasmopsis ciconiae
Mycoplasma anserisalpingitidis
Mycoplasmopsis anatis
Mycoplasmopsis synoviae
Mycoplasmopsis gallinacea
Mycoplasmopsis sturni
Mycoplasmopsis cricetuli
Mycoplasmopsis citelli
Mycoplasmopsis columboralis
Mycoplasma nasistruthionis
Mycoplasmopsis verecunda
Mycoplasma seminis
Mycoplasmopsis buteonis
Mycoplasmopsis glycophila
Mycoplasmopsis gallopavonis
Mycoplasmopsis mustelae
Mycoplasmopsis felis
Mycoplasmopsis leonicaptivi
Mycoplasmopsis bovirhinis
Mycoplasmopsis cynos
Mycoplasmopsis edwardii
Mycoplasmopsis canis
Mesomycoplasma mobile
"Candidatus Mycoplasma mahonii"
Mycoplasma todarodis
Mycoplasma marinum
Malacoplasma penetrans
Malacoplasma iowae
Ureaplasma canigenitalium
Ureaplasma zalophigenitalium
Ureaplasma miroungigenitalium
Ureaplasma diversum
Ureaplasma parvum
Ureaplasma urealyticum
Mycoplasmoides pneumoniae
Mycoplasmoides genitalium
Mycoplasmoides pirum
Mycoplasmoides alvi
Mycoplasmoides amphoriforme
Mycoplasmoides testudinis
Mycoplasma bradburyae
Mycoplasmoides imitans
Mycoplasma tullyi
Mycoplasmoides gallisepticum
Mycoplasmoides fastidiosum
"Candidatus Mycoplasma haematohominis"
"Candidatus Eperythrozoon haemobos"
Eperythrozoon canis
Eperythrozoon felis
Eperythrozoon parvum
Eperythrozoon suis
"Candidatus Eperythrozoon haemominutum"
"Candidatus Eperythrozoon haemolamae"
Eperythrozoon wenyonii
Eperythrozoon ovis
Acholeplasma laidlawii
Acholeplasma granularum
Acholeplasma oculi
Acholeplasma equirhinis
Acholeplasma hippikon
Acholeplasma equifetale

**Figure S1.** **Molecular phylogeny of the *Mycoplasmatales-Entomoplasmatales* clade.** The phylogenetic inference was based on a concatenated alignment of 30 conserved single-copy genes with 16,748 aligned amino acid sites. *Acholeplasma* was included as the outgroup. The maximum likelihood and the Bayesian trees have a symmetric difference value of 10 between their topologies. Those species with different phylogenetic placements are linked with horizontal lines between the two trees for comparison. Species included in the recent taxonomy revisions but have inconsistent phylogenetic placements are highlighted in red. Species not adhering to the recent taxonomy revisions are highlighted in blue (Hominis group) or orange (Pneumoniae group). Internal branches with bootstrap or posterior probability support below 80% have the exact support values labeled.
