## Supplementary material for "Delineating bacterial genera based on gene content analysis: a case study of the *Mycoplasmatales*–*Entomoplasmatales* clade within the class *Mollicutes*": Figure S3

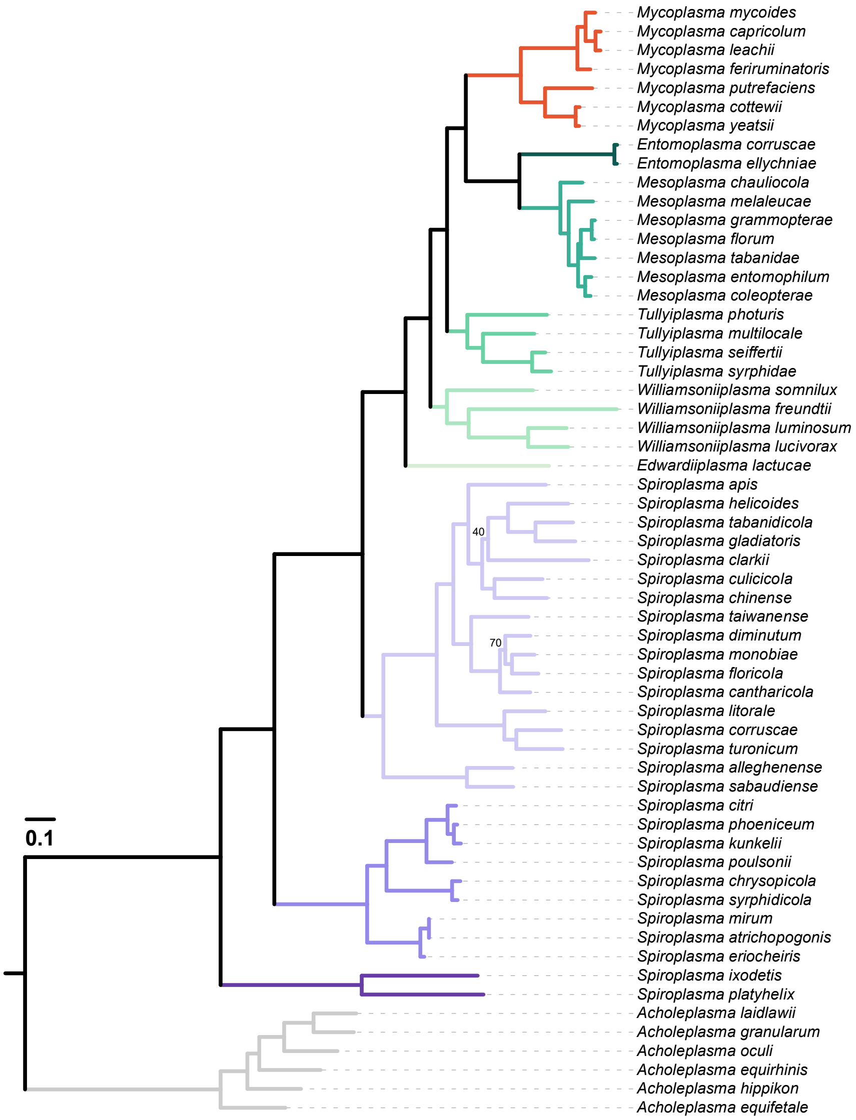


**Figure S3. Molecular phylogeny of the *Spiroplasma*-*Entomoplasmataceae*-Mycoides (SEM) group.** This maximum likelihood phylogeny included 53 ingroup species and 6 *Acholeplasma* spp. as the outgroup. The tree was inferred based on a concatenated alignment of 145 conserved single-copy genes with 62,530 aligned amino acid sites. Branches were color-coded according to the taxonomic assignments. Internal branches with bootstrap support below 80% have the exact support values labeled.
