## Supplementary material for "Delineating bacterial genera based on gene content analysis: a case study of the *Mycoplasmatales*–*Entomoplasmatales* clade within the class *Mollicutes*": Figure S4

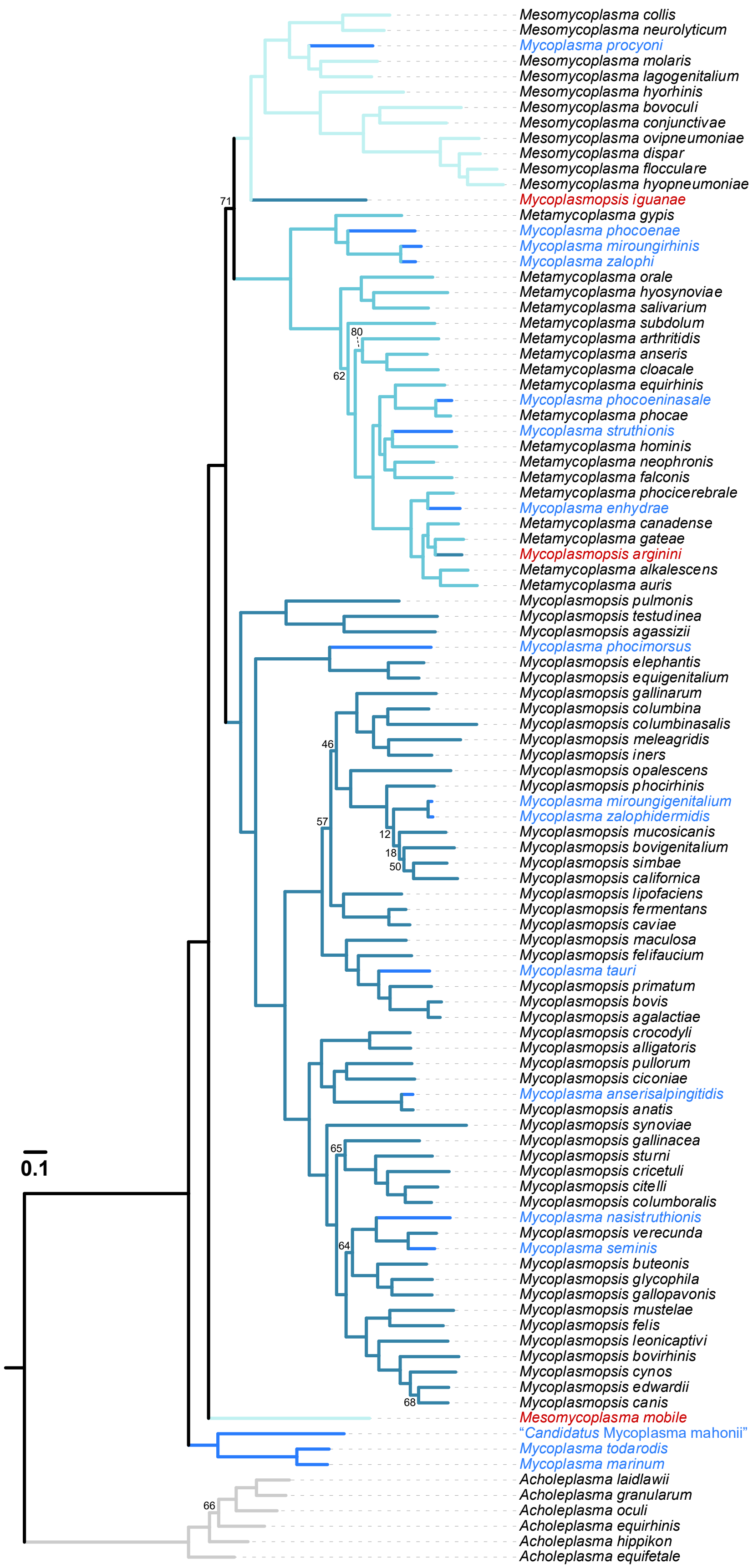
0.1
Mesomycoplasma collis
Mesomycoplasma neurolyticum
Mycoplasma procyoni
Mesomycoplasma molaris
Mesomycoplasma lagogenitalium
Mesomycoplasma hyorhinis
Mesomycoplasma bovoculi
Mesomycoplasma conjunctivae
Mesomycoplasma ovipneumoniae
Mesomycoplasma dispar
Mesomycoplasma flocculare
Mesomycoplasma hyopneumoniae
71
Mycoplasmopsis iguanae
Metamycoplasma gypis
Mycoplasma phocoenae
Mycoplasma miroungirhinis
Mycoplasma zalophi
Metamycoplasma orale
Metamycoplasma hyosynoviae
Metamycoplasma salivarium
Metamycoplasma subdolum
80
Metamycoplasma arthritidis
Metamycoplasma anseris
62
Metamycoplasma cloacale
Metamycoplasma equirhinis
Mycoplasma phocoeninasale
Metamycoplasma phocae
Mycoplasma struthionis
Metamycoplasma hominis
Metamycoplasma neophronis
Metamycoplasma falconis
Metamycoplasma phocicerebrale
Mycoplasma enhydrae
Metamycoplasma canadense
Metamycoplasma gateae
Mycoplasmopsis arginini
Metamycoplasma alkalescens
Metamycoplasma auris
Mycoplasmopsis pulmonis
Mycoplasmopsis testudinea
Mycoplasmopsis agassizii
Mycoplasma phocimorsus
Mycoplasmopsis elephantis
Mycoplasmopsis equigenitalium
Mycoplasmopsis gallinarum
Mycoplasmopsis columbina
Mycoplasmopsis columbinasalis
46
Mycoplasmopsis meleagridis
Mycoplasmopsis iners
Mycoplasmopsis opalescens
Mycoplasmopsis phocirhinis
Mycoplasma miroungigenitalium
57
Mycoplasma zalophidermidis
12
Mycoplasmopsis mucosicanis
18
Mycoplasmopsis bovigenitalium
50
Mycoplasmopsis simbae
Mycoplasmopsis californica
Mycoplasmopsis lipofaciens
Mycoplasmopsis fermentans
Mycoplasmopsis caviae
Mycoplasmopsis maculosa
Mycoplasmopsis felifaucium
Mycoplasma tauri
Mycoplasmopsis primatum
Mycoplasmopsis bovis
Mycoplasmopsis agalactiae
Mycoplasmopsis crocodyli
Mycoplasmopsis alligatoris
Mycoplasmopsis pullorum
Mycoplasmopsis ciconiae
Mycoplasma anserisalpingitidis
Mycoplasmopsis anatis
Mycoplasmopsis synoviae
65
Mycoplasmopsis gallinacea
Mycoplasmopsis sturni
Mycoplasmopsis cricetuli
Mycoplasmopsis citelli
Mycoplasmopsis columboralis
Mycoplasma nasistruthionis
Mycoplasmopsis verecunda
64
Mycoplasma seminis
Mycoplasmopsis buteonis
Mycoplasmopsis glycophila
Mycoplasmopsis gallopavonis
Mycoplasmopsis mustelae
Mycoplasmopsis felis
Mycoplasmopsis leonicaptivi
Mycoplasmopsis bovirhinis
Mycoplasmopsis cynos
Mycoplasmopsis edwardii
68
Mycoplasmopsis canis
Mesomycoplasma mobile
“Candidatus Mycoplasma mahonii”
Mycoplasma todarodis
Mycoplasma marinum
Acholeplasma laidlawii
Acholeplasma granularum
66
Acholeplasma oculi
Acholeplasma equirhinis
Acholeplasma hippikon
Acholeplasma equifetale

**Figure S4. Molecular phylogeny of the Hominis group.** This maximum likelihood phylogeny included 95 ingroup species and 6 *Acholeplasma* spp. as the outgroup. The tree was inferred based on a concatenated alignment of 73 conserved single-copy genes with 34,935 aligned amino acid sites. Branches were color-coded according to the taxonomic assignments. Internal branches with bootstrap support below 80% have the exact support values labeled.
