## Supplementary material for "Delineating bacterial genera based on gene content analysis: a case study of the *Mycoplasmatales*–*Entomoplasmatales* clade within the class *Mollicutes*": Figure S5

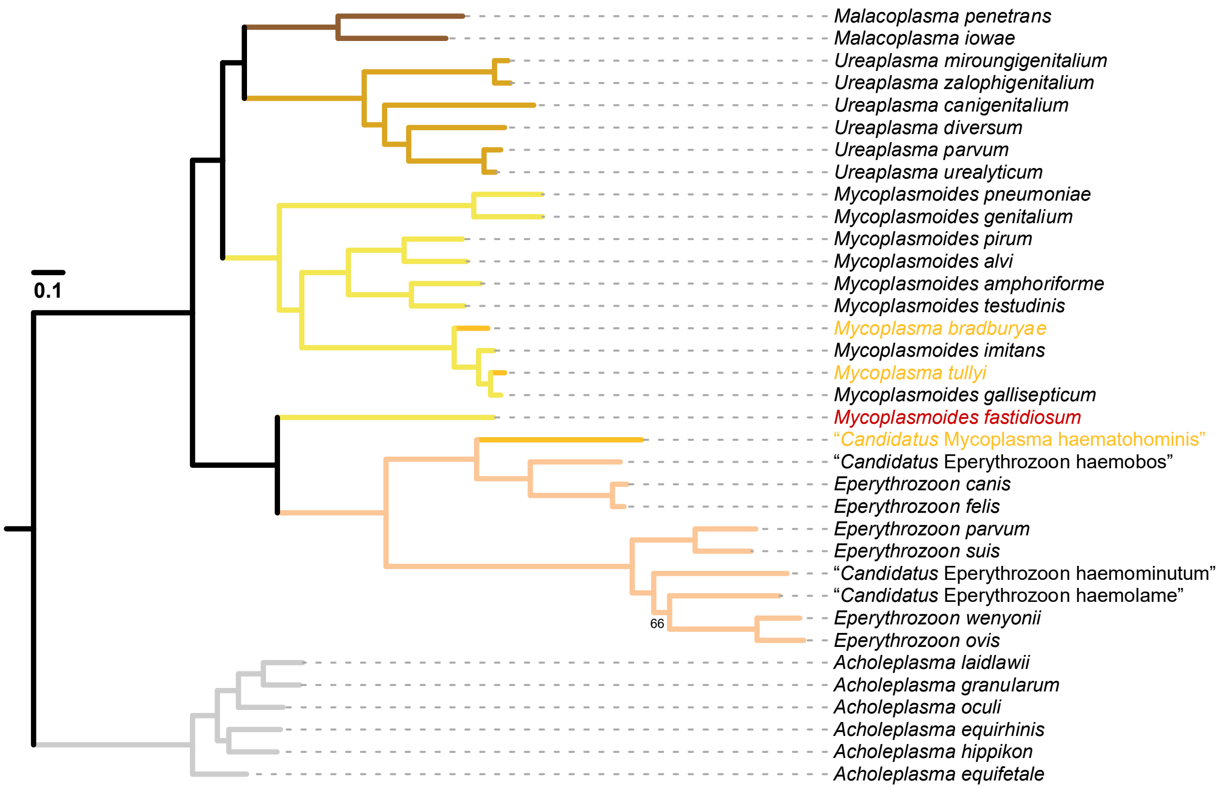


**Figure S5. Molecular phylogeny of the Pneumoniae group.** This maximum likelihood phylogeny included 29 ingroup species and 6 *Acholeplasma* spp. as the outgroup. The tree was inferred based on a concatenated alignment of 81 conserved single-copy genes with 37,270 aligned amino acid sites. Branches were color-coded according to the taxonomic assignments. Internal branches with bootstrap support below 80% have the exact support values labeled.
