## Supplementary figures and images for "Delineating bacterial genera based on gene content analysis: a case study of the *Mycoplasmatales*–*Entomoplasmatales* clade within the class *Mollicutes*"

### Figure S6

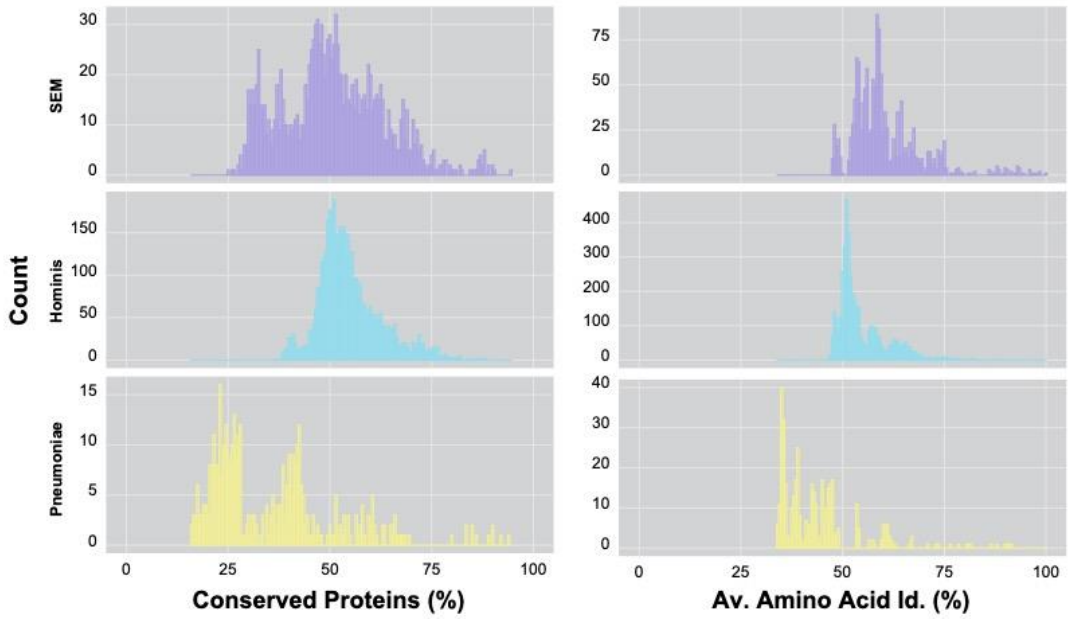


**Figure S6. Frequency distribution of pairwise genome similarities.**
